# The Neuromuscular system of *Chironomus vitellinus* (Diptera: Chironomidae)

**DOI:** 10.1101/2025.06.01.657287

**Authors:** Roberto Reyes-Maldonado, Alonso Ramírez, Bruno Marie

## Abstract

Chironomids are important laboratory model organisms used to assess toxicity in freshwater environments. Cell and tissue features are not commonly used as chironomid markers to detect toxicity, but they could be extremely helpful in identifying acute and chronic effects of pollutants. The nervous system is an excellent cellular candidate since it is reactive to toxic substances. However, a detailed description of the chironomid nervous system is required prior to considering it as a candidate for a cellular toxicity marker. The present study describes the central ganglia, nerves, axons, and the neuromuscular system of *Chironomus vitellinus* (Freeman, 1961) to facilitate its use as a model organism in environmental studies. We find that the structure of the *C. vitellinus* central nervous system is identical to that observed in other *Chironomus* larvae. We then focused our study on the first abdominal segment and labeled the 31 hemi-segmental muscles according to a nomenclature based on their position and orientation. We also characterized their innervation and assigned the nerves a nomenclature based on their terminals’ location in the muscle tissue. Finally, we investigated the neuromuscular junctions (NMJs) throughout this segment and defined four types of NMJs illustrating their great variability in size and shape. We selected a model NMJ, VEL 2, and quantified its mean bouton number and muscle size. Together with documenting a neurobiological system that could be informative to insects’ comparative biology, these results could help establish the *Chironomus* NMJ as an aquatic toxicity marker.

## Introduction

The aquatic larvae of various species of chironomids (Diptera: Chironomidae) have become a model of choice for assessing freshwater contaminants. Indeed, their larvae are easy to maintain under laboratory conditions, resist environmental changes, feed and burrow in benthic sediments, respond to many pollutants, and have a short life cycle [1]. When assessing toxicity, a series of markers have been used, from biochemical and molecular markers (BMMs) [2,3] to morphological and fitness markers (MFMs) [1].

The main benefit of using BMMs to assess toxicity is that these kinds of markers show the effects of chemicals before adverse effects are detectable at higher levels of biological organization [4]. Also, BMMs can provide important insights regarding how organisms deal with toxic chemicals and the mechanisms of toxicity [5]. In the case of MFMs, the main benefit is that the effects observed on individuals can be used to anticipate ecological effects. Despite the benefits offered by these markers, there are disadvantages. For example, BMMs are expensive to use in terms of the equipment needed and running costs [5] while MFMs are negatively influenced by the animal behavior and capacity for producing detoxification mechanisms [4].

Although BMMs and MFMs provide important information when assessing toxicity in freshwater environments, cellular and tissue markers (CTMs) represent a better alternative [6]. These markers provide information on the integration of changes produced at molecular, biochemical, and physiological levels that have resulted from chemical exposure [7]. As a result, contaminant-induced cellular and tissue alterations can be related to the health and fitness of individuals allowing further extrapolation to population or community effects [7]. When compared to BMMs and MFMs, CTMs present a more cost-effective way to verify toxicant effects and present better responses to long-term alterations with higher ecological relevance [6].

Although a useful alternative, CTMs are rarely used as markers for assessing toxicity due to the lack of histological descriptions in this insect group. Having noticed this gap in knowledge, recent studies have provided new anatomical descriptions using modern histological techniques to identify new cellular and tissue markers (CTMs) for the assessment of freshwater contaminants [6,8]. Nevertheless, these studies do not focus on the use of the nervous tissue, a tissue that should be considered to assess toxicity given that contaminants can affect its function [9,10] and development [11,12].

The nervous system, as studied in the *Drosophila melanogaster* model, could be used as reference for the implementation of a quantifiable cellular marker for a *Chironomus* species. Because they are easily accessible, documenting neuromuscular junctions (NMJs, [13,14]) could provide a way to quantify and assess the toxicity of the animal milieu. *Chironomus* larval musculature and nervous system have been described separately [6,8,15]. Nevertheless, the general description of the larval musculature in its entirety has not been presented and no terminology allowing the identification of the different muscles has been defined [15]. Similarly, the central nervous system and its main projections to the periphery have only been described showing the different ganglia [6,8,15] and their corresponding main nerves [15]. Other studies have focused on describing the ultrastructure of the neural sheath, glial cells, and neurons, while others have identified the location of selected transmitters, transmitter related enzymes and neuropeptides through the ganglia and alimentary canal [16–18]. No study has provided a complete description of the neuromuscular system characterizing and naming the larval muscles and the nerve terminals innervating them.

NMJs, which are responsible for inducing muscle contraction and the animal’s locomotion [19], have been extensively studied in fruit flies [20], crustaceans [21], frogs [22], and mice [23]. In this study, we used immunohistochemical synaptic markers to describe in detail the neuromuscular anatomy (muscles, nerves and NMJs) of the first abdominal segment in the larvae of the broadly distributed chironomid *Chironomus vitellinus.* In addition, we selected a model NMJ to quantify features such as synaptic size. We assessed synaptic size by quantifying the number of synaptic boutons, the varicosities present at each terminal and containing the neurotransmitter release apparatus [24]. We hypothesize that our detailed descriptions of the neuromuscular system in chironomids will establish a framework for the study of the *Chironomus* NMJ as a CTM for future toxicological studies.

## Materials and Methods

### Animal husbandry

This study used *C. vitellinus* as a model chironomid species. Previously, this species was referred to as *Chironomus* sp. ‘Florida’ in Reyes-Maldonado et al., 2021 [25] before being recently identified as *C. vitellinus* [26,27]. *C. vitellinus* has a widespread distribution, with reports from Africa, Asia, Oceania, and, more recently, the Americas [26]. Based on our research, these chironomids can be easily collected in both urban and rural areas of Puerto Rico, particularly in natural temporary pools and animal waterers.

The larvae used in this study were acquired from laboratory-reared colonies at the Institute of Neurobiology, University of Puerto Rico. These colonies were obtained from egg masses collected in the field using a previously described methodology [25]. The larvae studied were fed with TetraMin Tropical Flakes Fish Food provided as a slurry at 2 mg / individual / day and maintained at 27°C with a photoperiod of 12:12 light/dark. The 4th larval, and final, instar was the stage used for all the descriptions presented in this study. They were identified by their size (9-14 mm approximately) and bright red coloration (Figure 1A). Animals of both sexes were used in these procedures.

**FIGURE 1:**
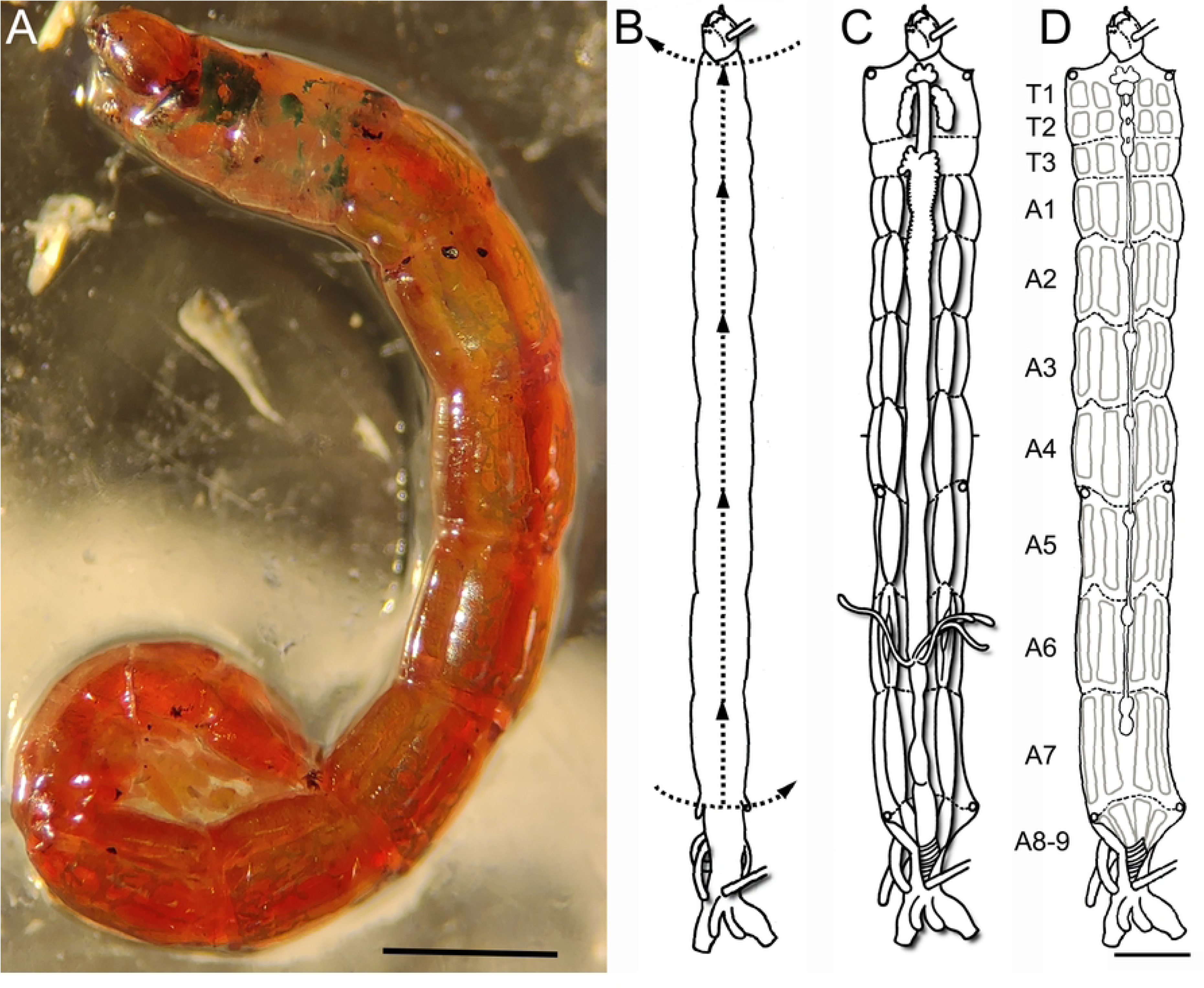
The fourth larval instar of *Chironomus vitellinus* and the dissection technique used to obtain neuromuscular tissue. **(A)** The entire body of the 4^th^ instar larvae is bright red with some green spots in the thorax in completely mature larvae. This stage is characterized by individuals having a length between 9-12mm. **(B-D)** Schematic view of the dissection process designed for studying the neuromuscular tissue of *C. vitellinus*. **(B)** Initial position of larvae in the plate after pinning the larvae at its anterior and posterior extremities. The dashed lines and arrows represent the initial cuts of the dissection process. **(C)** Stretching the open larvae reveals extensive fat tissue, digestive tract, salivary glands, and tracheal system. **(D)** After removing the visceral tissue and fat, we can observe the segmental organization (T1 to A9) of both the CNS (chain of ganglia within the median plane) and the muscles (light grey rectangles). Scale: 1mm.

### Dissection

Tissue was dissected following an adapted methodology described by Sanhueza et al., 2016 [28]. Briefly, 4th instar larvae were attached, dorsal face up, to a Sylgard dish by pinning the space between the head and first abdominal segment and the last abdominal segment. Subsequently the larvae were covered with HL3 Ringer’s saline solution (70mM NaCl_2_, 4mM KCl, 43mM MgCl_2_, 10 mM NaHCO_3_, 115mM Sucrose, 5mM HEPES) and dissected using microdissection scissors under the stereoscope. The first incision was made transversally above the heart at abdominal segment 7/8. A second incision was made intersecting the first, proceeding anteriorly following the aorta until reaching the head of the animal. A third incision was made transversally at the end of the second incision to open the thoracic cavity (Figure 1B). Four pairs of pins were placed at each border of the 1st thoracic segment, 1st abdominal segment, 5th abdominal segment and the 7th abdominal segment in order to open the thoracic and abdominal cavities by stretching the tissue (Figure 1C, D).

### Muscle characterization and imaging

Dissected tissue was stained using bromophenol blue (Millipore) diluted in Ringer’s saline solution (5mg/ml) for 3 minutes. Muscle layers were successively removed starting with the most internal (visceral) muscles and finishing with the most external (cuticular) muscles. Within each layer, muscle arrangements were described by drawing under a stereomicroscope. Additional stained tissue was mounted on microscope slides, observed under an AmScope T490B-DK microscope, and photographed with a piece of metric paper (1mm grid) using a 108MP phone camera (Motorola Edge, 2021) attached to the ocular. A scale was added to the obtained images in ImageJ (National Institutes of Health; http://imagej.nih.gov/ij/) by transforming pixel units to the known length of the metric paper grid. These images were used to produce more detailed drawings using the GNU Image Manipulation Program (GIMP 2.10.10, www.gimp.org). We defined a nomenclature based on position and orientation to label the muscles of the ventral and lateral region similar to previous work carried out on *Drosophila* larval anatomy [29,30].

### Nervous tissue immunohistological characterization and imaging

After dissection, the stretched tissue was fixed with Bouin’s solution (Sigma) for 1 minute and washed three times with PBT solution [1x Phosphate-buffered Saline (PBS) and 0.1% Triton-X 100]. Guts and body fat were then carefully removed with tweezers and a bent pin head. After removing the pins, the tissue was washed and permeabilized in five PBT baths for an hour (12 min/bath).

Immunohistochemistry was used to visualize neuronal tissue. We assessed the cross reactivity of antibodies raised against *Drosophila melanogaster* proteins in the *C. vitellinus* tissue (Table 1). The goat Cy3-conjugated anti-HRP antibody (1:300 in PBT; Jackson ImmunoResearch; [31] and the following primary antibodies: mouse anti-Syn (1:20 in PBT; Developmental Studies Hybridoma Bank (DSHB) [32]), mouse anti-Dlg (1:20 in PBT; DSHB [33]), rabbit anti-Dlg (a gift from Dr Budnik; 1:150 in PBT[34]), mouse anti-FAS2 (1:10 in PBT; DSHB[35]), mouse anti-Futsch (1:10 in PBT; DSHB [36]), mouse anti-α -Tubulin (1:3000 in PBT; DSHB; [37,38]), mouse anti-Bruchpilot (1:50 in PBT; DSHB; [39]), mouse anti-Synaptotagmin (1:50 in PBT; DSHB;[40]), mouse anti-GluRIIA (1:20 in PBT; DSHB; [41,42]) and rabbit anti-GluRIII (a gift from Dr DiAntonio; 1:2000 in PBT; [43,44]) were tested. The immunohistochemistry process was initiated by incubating clean *C. vitellinus* tissue in the diluted primary antibody overnight at 4°C. This incubation was followed by five PBT washes during 1hr (12min/wash) and a second 1hr incubation of the tissue was carried out in the dark with the diluted secondary antibody [Alexa Fluor 488-conjugated AffiniPure goat anti-mouse or anti-rabbit IgG (1:300; Jackson ImmunoResearch) and the Cy3 conjugated anti-HRP. A second wash of five changes of PBT in the dark was applied and the preparations were mounted on glass microscope slides using Vectashield (Vector Labs) as mounting media. The preparations were sealed with nail polish and stored in a cool dark place until analysis.

**TABLE 1:** Testing the chironomids NMJ immunoreactivity with antibodies used to study the *Drosophila* NMJ. 11 antibodies were tested against the *C. vitellinus* preparations. Unless specified, monoclonal antibodies were used to assess *C. vitellinus* tissue immunoreactivity. 7 such antibodies revealed some immunoreactivity in *C. vitellinus* consistent with the role of the *Drosophila* proteins they were raised against. These antibodies revealed synaptic structures such as neuronal membranes, cell adhesion molecules, synaptic boutons, pre-synaptic active zones, and post-synaptic differentiation.

| Antibody | Target | Tissue visualized | References | *Chironomus*<br>Immunoreactivity |
| --- | --- | --- | --- | --- |
| Anti-HRP | Neuronal cell surface proteins | Presynaptic membrane | [31] | Yes |
| Anti-FAS2 | Cell adhesion molecule Fas 2 | Synapse | [35] | Yes |
| Anti-α-Tubulin | α-Tubulin isoform in microtubules | Cytoskeleton | [37,38] | Yes |
| Anti-Syn | Synaptic vesicle associated protein Synapsin | Synaptic boutons | [32] | Yes |
| Anti-Brp | Active zones marker Bruchpilot | Presynaptic active zones | [39] | Yes |
| Anti-DLG<br>(polyclonal) | Postsynaptic scaffold protein discs large | Postsynaptic densities | [34] | Yes |
| Anti-DLG<br>(monoclonal) | Postsynaptic scaffold protein discs large | Postsynaptic densities | [33] | No |
| Anti-GluR3 | Glutamate receptor subunit 3 | Postsynaptic densities | [43, 44] | No |
| Anti-GluR2A | Glutamate receptor subunit 2A | Postsynaptic densities | [41, 42] | No |
| Anti-Syt1 | Synaptic vesicle associated protein Synaptotagmin 1 | Synaptic boutons | [40] | No |
| Anti-Fustch | Microtubule associated protein MAP1B | Cytoskeleton | [36] | No |

The specimens were scanned using a Nikon Eclipse Ti inverted A1R laser scanning confocal microscope. The NIS elements Advance Research 4.5 acquisition and analysis software was used for image acquisition. Complete body tissue recordings were obtained over the Z-axis (0.5 µm depths) at 20X magnification. Recording of specific features such as ganglia and nerves were obtained at 40X (0.2 µm depths) while the NMJs at 40X and 100X (0.2 and 0.1 µm depths, respectively). Large structures were recorded by scanning multiple areas in the X or Y axis with an overlap of 20% of the scanned area. Using Image J, images were produced as Maximum Intensity Projections over the Z-axis of the stack of images. Figures were made using Adobe Photoshop 2023. In Figure 5, to better visualize some NMJ structures, the background of the image was attenuated by using the GNU Image Manipulation Program (GIMP; https://www.gimp.org/). Briefly, the silhouette of the studied synaptic terminal was selected by hand on the color channel containing the HRP staining. The width of this selection was increased to 1 pixel size and later inverted to erase the area of no interest. GIMP was also used to create diagrams from raw images to show the location and appearance of ganglia, nerves and NMJs. A nomenclature to these structures was assigned based on previous work [15,29,30].

### Characterization of a model NMJ

The selected model NMJ was observed in 10 larval preparations (19 NMJs) immunolabeled with the anti-Syn and anti-HRP antibodies. The bouton numbers were quantified under an Eclipse 80i microscope at 60X. We quantified the mean number and the variability of the samples using “GraphPad Prism 6”. This software was also used to carry out the Shapiro-Wilk normality test to assess the distribution of the samples.

## RESULTS

### Antibody reactivity in the nervous and muscle tissue of C. vitellinus

To investigate the different anatomical features of the *C. vitellinus* larvae, we decided to rely on well characterized antibodies in another Diptera, *Drosophila melanogaster*. We tested eleven antibodies routinely used to characterize neurons and their synapses in *Drosophila*. Six of these antibodies presented immunoreactivity in *C. vitellinus* consistent with the described labeling of *Drosophila* tissue (Table 1). With these antibodies we were able to visualize neuronal membranes (anti-HRP; [31]), synaptic vesicles (anti-Syn; [32]), neuronal cell adhesion molecule (anti-FAS2; [35]), microtubules (anti-α-Tubulin; [37]), active zones (anti-Brp; [39]), and post-synaptic scaffold proteins (anti-Dlg; [34]). In our study we used the neuronal membrane immunoreactivity revealed by the anti-HRP antibody to visualize the ganglia, nerves, and axons. The anti-Syn and anti-HRP antibodies were used to reveal the neuromuscular system. Like in *Drosophila*, synaptic boutons were visualized and quantified using the anti-Syn antibody.

### Accessing the C. vitellinus tissue and observation of the basic larval structure

We dissected the 4^th^ Instar larvae of *C. vitellinus* (Figure 1A). Upon dissection (described in methods; Figure 1B), the first structures exposed are the digestive tract and the salivary glands (Figure 1C). Once these structures are removed, an inner fatty layer can be observed covering the muscles and the central nervous system from the first to the last abdominal segment. This fatty layer is segmentally organized and reduces the visibility of the neuromuscular tissue especially around the central nervous system and the lateral muscles. Although this fatty layer cannot be eliminated, its careful partial removal in most of the areas of the preparation is required to visualize the neuromuscular system. The three thoracic and eight of the nine abdominal segments of the larva provide easily identifiable muscles, nerves and NMJs (schematic diagram; Figure 1D). While each thoracic segment (T1 to T3) presents a unique muscular organization, the arrangement of muscles is almost identical in the first seven abdominal segments (A1 to A7) and differs in the eighth and ninth abdominal segments. The muscles within different segments are innervated by a central nervous system (CNS) that is segmented and organized as a series of ganglia. We therefore decided to describe the CNS in more detail.

### The central nervous system

Although the principal anatomical CNS features had already been described in other species within the *Chironomus* group [6,8,15], it was important to determine whether the anatomy of the *C. vitellinus* CNS was identical.

The *C. vitellinus* CNS is composed of a brain (also called supraoesophageal ganglion), a suboesophageal ganglion, three thoracic ganglia, and eight abdominal ganglia (Figure 2). The brain (br.) is located out of the head in the prothorax, and it is the largest and most prominent of the ganglia. Lying beneath the brain and in the same thoracic segment, the suboesophageal ganglion (s.oes.g.) and the prothoracic ganglion (pro.g.) stands out. The mesothoracic ganglion (mes.g.) and metathoracic ganglion (met.g.) are located in the mesothorax. The first abdominal ganglion (1 ab.g.) is in the metathorax, leaving the first abdominal cavity unoccupied. Abdominal ganglia 2-6 (2-6 ab.g.) occupy their respective abdominal segments. The 7^th^ and 8^th^ abdominal ganglia are both located in the 7^th^ abdominal segment forming the terminal ganglion of the nerve cord (t.g.).

**FIGURE 2:**
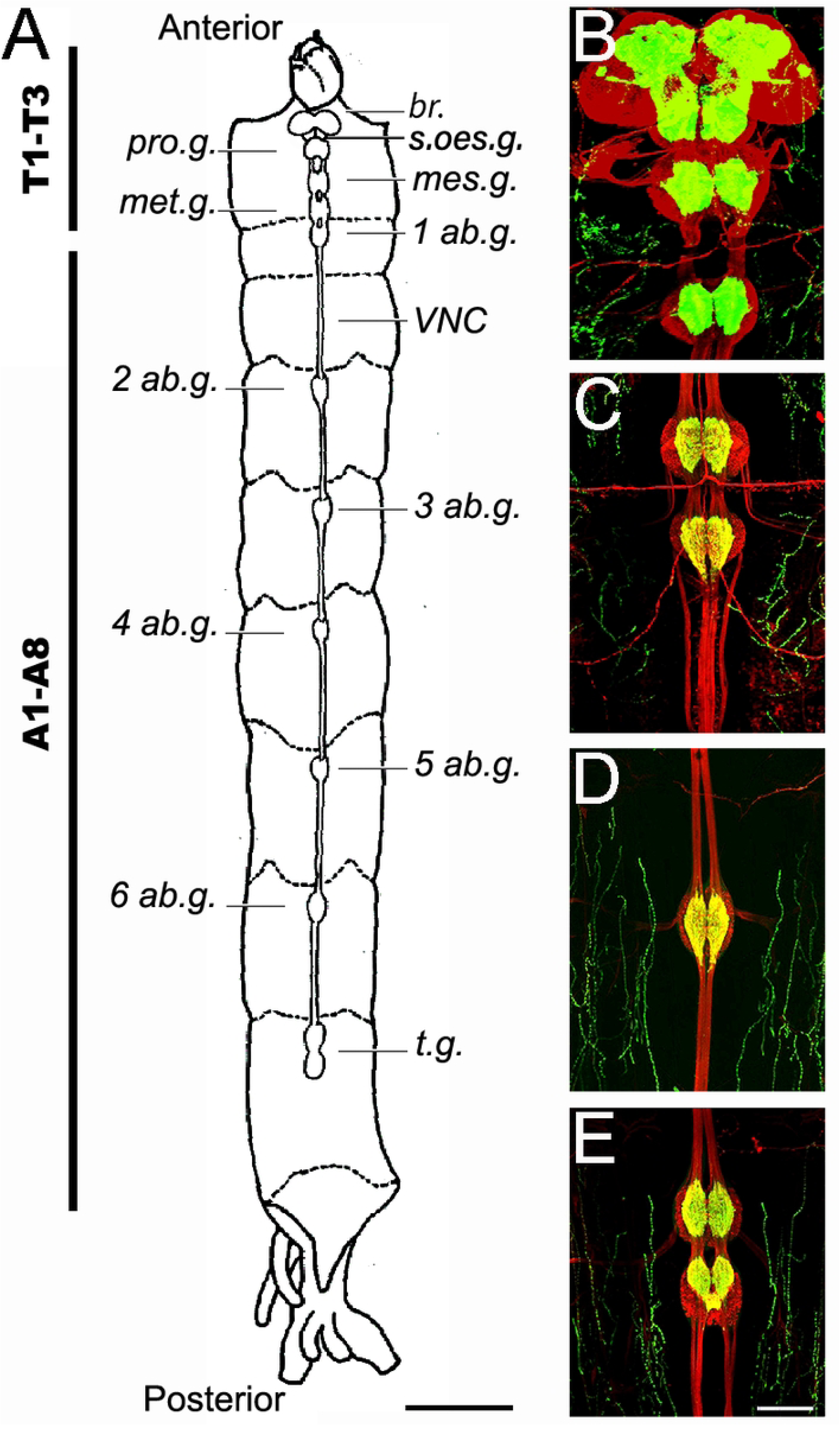
Principal components of the central nervous system of the *Chironomus vitellinus* larva. **(A)** General schematic view of the larval CNS with nomenclature of principal structures. Scale: 1 mm **(B-D)** Anti-HRP (red) and anti-Synapsin (green) immunofluorescence reveal the different CNS structures. Scale bar: 100um. **(B)** Representative confocal photograph of the brain (supraoesophageal ganglion), suboesophageal ganglion, prothoracic ganglion and mesothoracic ganglion **(C)** Representative confocal photograph of the metathoracic ganglion and the first abdominal ganglion **(D)** Representative confocal photograph of an abdominal ganglion from the 5th abdominal ganglion **(E)** Representative confocal photograph of terminal ganglia. Abbreviations: Tx: Thoracic segment x; Ax: Abdominal segment x; br.: Brain; s.oes.g. suboesophageal ganglion; pro.g.: prothoracic ganglion; mes.g.: mesothoracic ganglion; met.g.: metathoracic ganglion; xAb.g.: abdominal ganglion x; t.g.: terminal ganglion; VNC: ventral nerve cord.

Nerves exit each ganglion to their respective region: nerves from the supra and suboesophageal ganglia reach only the head, the ones from the thoracic ganglia reach the thorax, and the ones from the abdominal ganglia reach their respective abdominal segments. Each ganglion is composed of two lobes and is interconnected to other ganglia through a series of nerve fibers, forming the ventral nerve cord (*VNC*).

Our results show that indeed the anatomical organization of the *C. vitellinus* CNS is identical to that observed in other *Chironomus* larvae such as *C. dorsalis* [15], *C. sancticaroli* [6], and *C. columbiensis* [8].

### The musculature of the first abdominal segment

After describing the global larval architecture for both muscles and CNS, we decided to detail the organization of one segment. The first abdominal segment seemed the perfect candidate since it is easily accessible due to its central location and to the fact that the ganglion innervating its different muscles is located anterior to the segment itself (Figure 3). This last characteristic not only ensures unaltered visual access to the muscles, but it also facilitates potential future electrophysiological studies since the increased distance between muscle and ganglion also means longer and more accessible nerves.

**FIGURE 3:**
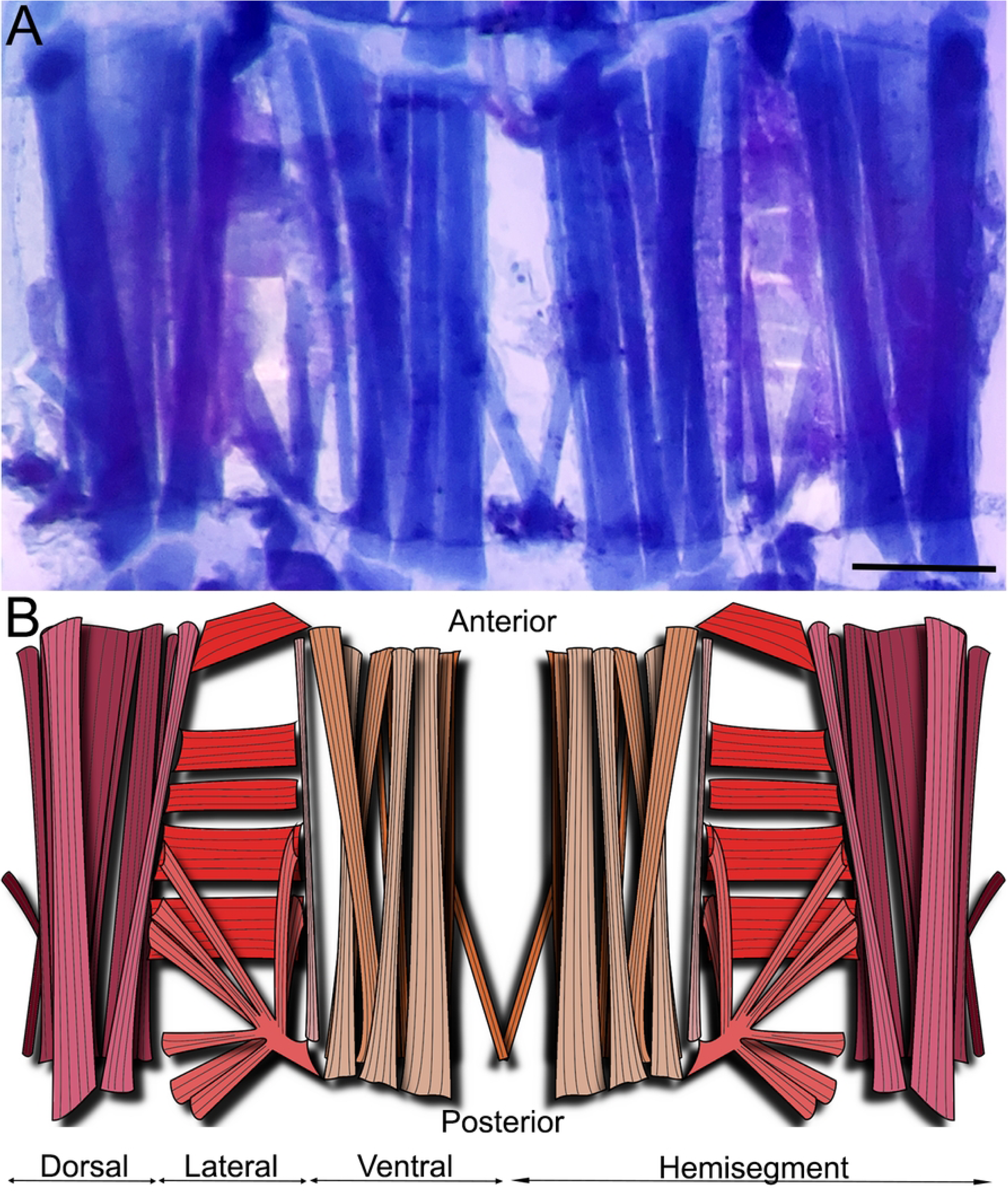
Musculature of the first abdominal segment of *Chironomus vitellinus*. **(A)** First abdominal segment as observed with the bromophenol blue staining. Scale is 200 µm. **(B)** Diagram showing the position and orientation of muscles in the first abdominal segment. Purple: dorsal muscles; Red: lateral muscles; Orange: ventral muscles. Light colors represent visceral/interior muscles while dark colors cuticular/exterior ones.

Three orientations of muscles (longitudinal, transverse, and oblique) can be distinguished in the ventral, lateral and dorsal area of the thoracic and abdominal segments (Figure 3). The first abdominal hemisegment presents 31 muscles: 12 lateral, 10 ventral and 9 dorsal muscles. Regarding the orientation of the muscles in the ventral and lateral area, we can observe 8 oblique, 9 longitudinal and 5 transverse muscles per hemisegment. Muscle fibers are present in two layers, presenting 3 main muscles in the internal part and 7 muscles in the external part of the ventral region. The following five groups of muscles were defined: ventral external longitudinal (VEL), ventral internal longitudinal (VIL), ventral oblique (VO), lateral transverse (LT), and Lateral oblique (LO). The distribution of these muscles is described in detail in Figure 4. Figure 4C describes the external muscle layers and illustrates the nomenclature we used to identify the muscles namely SBM, VEL 1 to 4, VEL 6, VO1, LT 1 to 4, LO1, LO 4 to 9. Underneath this sheet of muscles, we can find the internal muscles (VIL 1 to 3, VEL 5; Fig 4D). As depicted in Figure 4, NMJs are visible and identifiable on every muscle. We therefore underwent the process of naming the nerves innervating the muscular architecture.

**FIGURE 4:**
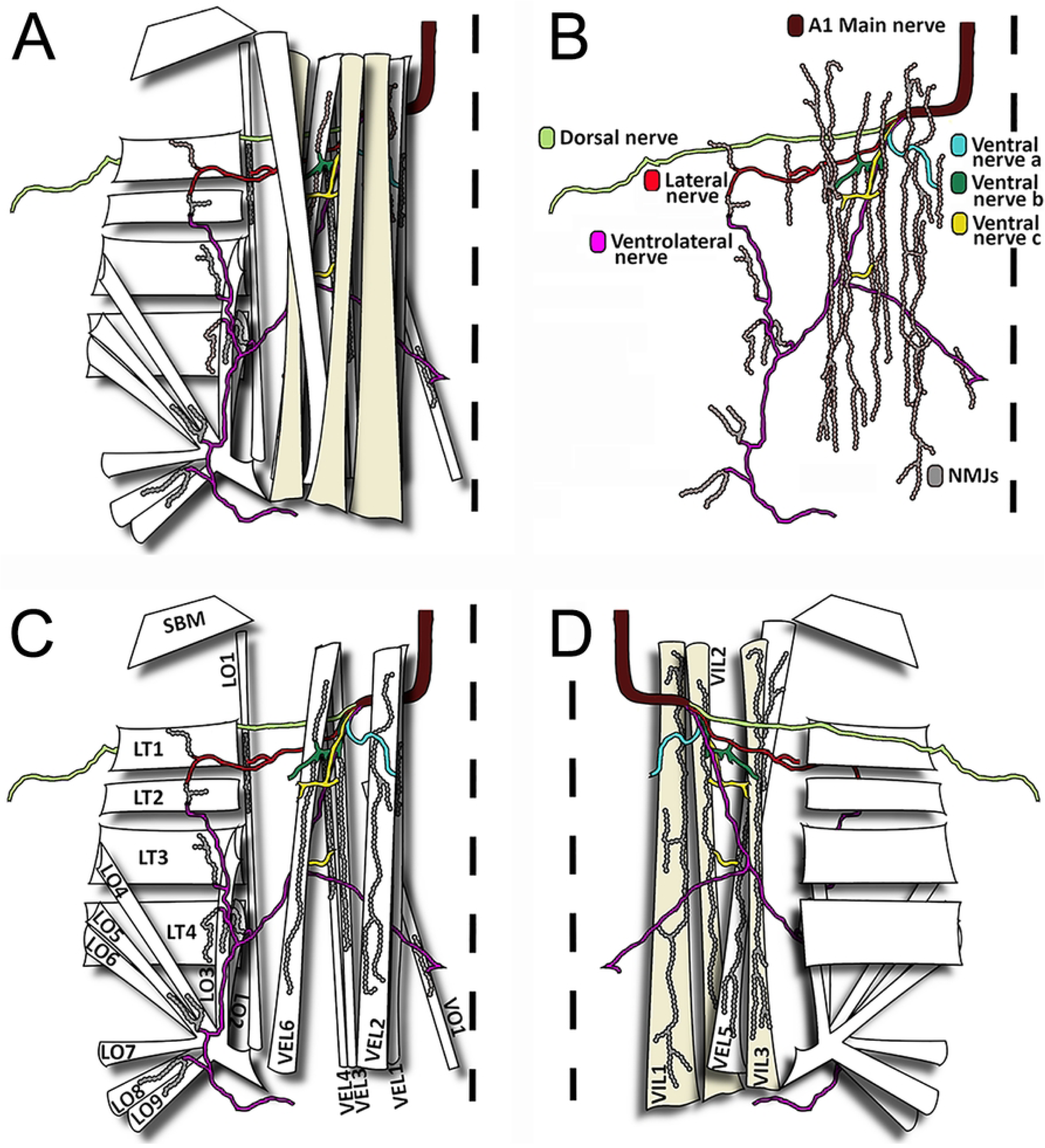
The neuromuscular organization of the first abdominal segment’s lateral and ventral regions. **(A)** Schematic representation of body wall muscles of one hemisegment with their respective peripheral nerve branches and neuromuscular junctions. Note that there are different layers of muscles and innervation going from the most cuticular/exterior to the more visceral/interior **(B)** Representation of the NMJs with their respective nerves in a view in which muscles were removed. Individual nerves are named and NMJs are drawn **(C)** View of the cuticular / exterior layer of muscles and their innervation. Individual muscles are named **(D)** View of the visceral / Interior layer of muscles and their innervation shows the morphology of typical NMJs. **(A-D)** The midline, that separates the two bilaterally symmetrical hemisegments, is represented by a dotted line. Each nerve branch is represented with a single color to show target muscles. Muscle and nerve nomenclature are based on position, orientation, and number (Bate 1993 and Landgraf et al. 1997). Abbreviation: Position V: ventral; VI: Ventral internal; VE: Ventral external L: lateral. Orientation: L: longitudinal O: oblique; T: transverse, SBM, segment border muscle.

### Nerves projecting to the first abdominal segment

We named “main nerves’’ the two hemisegmental nerves originating from the first abdominal ganglia. There are six nerve tracts originating from the main nerve. We assigned these previously unnamed nerve tracts a nomenclature based on their nerve terminals’ location in the muscle tissue in a manner reminiscent of previously published work in *Drosophila* [13,30]. The six nerve tracts are: ventral nerve a, ventral nerve b, ventral nerve c, ventrolateral nerve, lateral nerve, and dorsal nerve (Figure 4B). The synaptic terminals of the *ventral nerve a, b* and *c* only reach ventral muscles; the synaptic terminals of the *ventrolateral nerve* reach a single ventral muscle but also few lateral muscles; the synaptic terminals of the *lateral nerve* reach only lateral muscles; and the synaptic terminals of the *dorsal nerve* reach all the dorsal muscles (Figure 4).

### The first abdominal segment NMJs

We used the anti-Synapsin and the anti-HRP antibody labeling to observe the location of the NMJs in the *C. vitellinus* tissue. Because Synapsin is a synaptic vesicle protein [32], the anti-Synapsin fluorescence concentrates in the synaptic boutons of the NMJ. Using this immunofluorescence, we noticed that some synapses showed clearly separated large-sized boutons with a round morphology, well distributed through the NMJ. Thanks to these characteristics, we were able to distinguish individual synaptic boutons, giving us the opportunity to quantify the number of boutons per synapse (for example Figure 5A and 6A). Any NMJ in which the anti-Synapsin immunoreactivity labeling lacked this description was cataloged as unquantifiable (for example Figure 5B and 6B). The anti-HRP antibody was used to visualize the neuronal membrane (axon, boutons, and inter-bouton synaptic area) and allowed the delineation of the NMJ and the identification of its origin. Integrating both antibodies, we determined the location of each NMJ and the axon from which it is originating.

**FIGURE 5:**
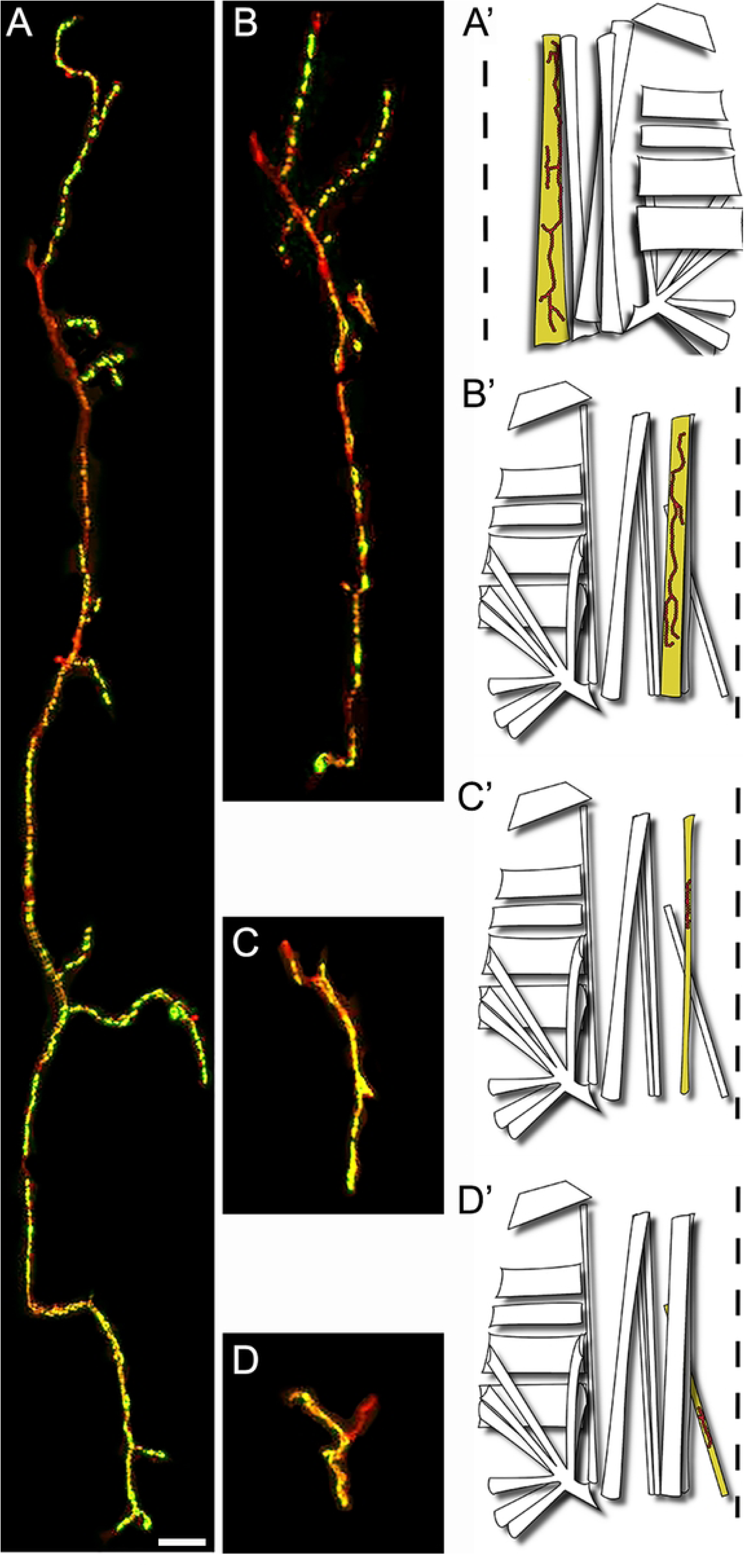
NMJs Diversity in the first abdominal segment. **(A)** Representative confocal photograph of a large ventral interior NMJ VIL1 revealed by anti-HRP (red) and anti-Synapsin (green) immunoreactivity. **(A’)** Diagram showing the location of the VIL1 muscle and its NMJ within the hemisegment **(B)** Representative confocal photograph and **(B’)** diagram of a medium ventral external NMJ on muscle VEL2 **(C)** Representative confocal photograph and **(C’)** diagram of a small NMJ on muscle LO1 **(D)** Representative confocal photograph and **(D’)** diagram of a very small NMJ on muscle VO1. Scale bar: 100 µm (A-D)

**FIGURE 6:**
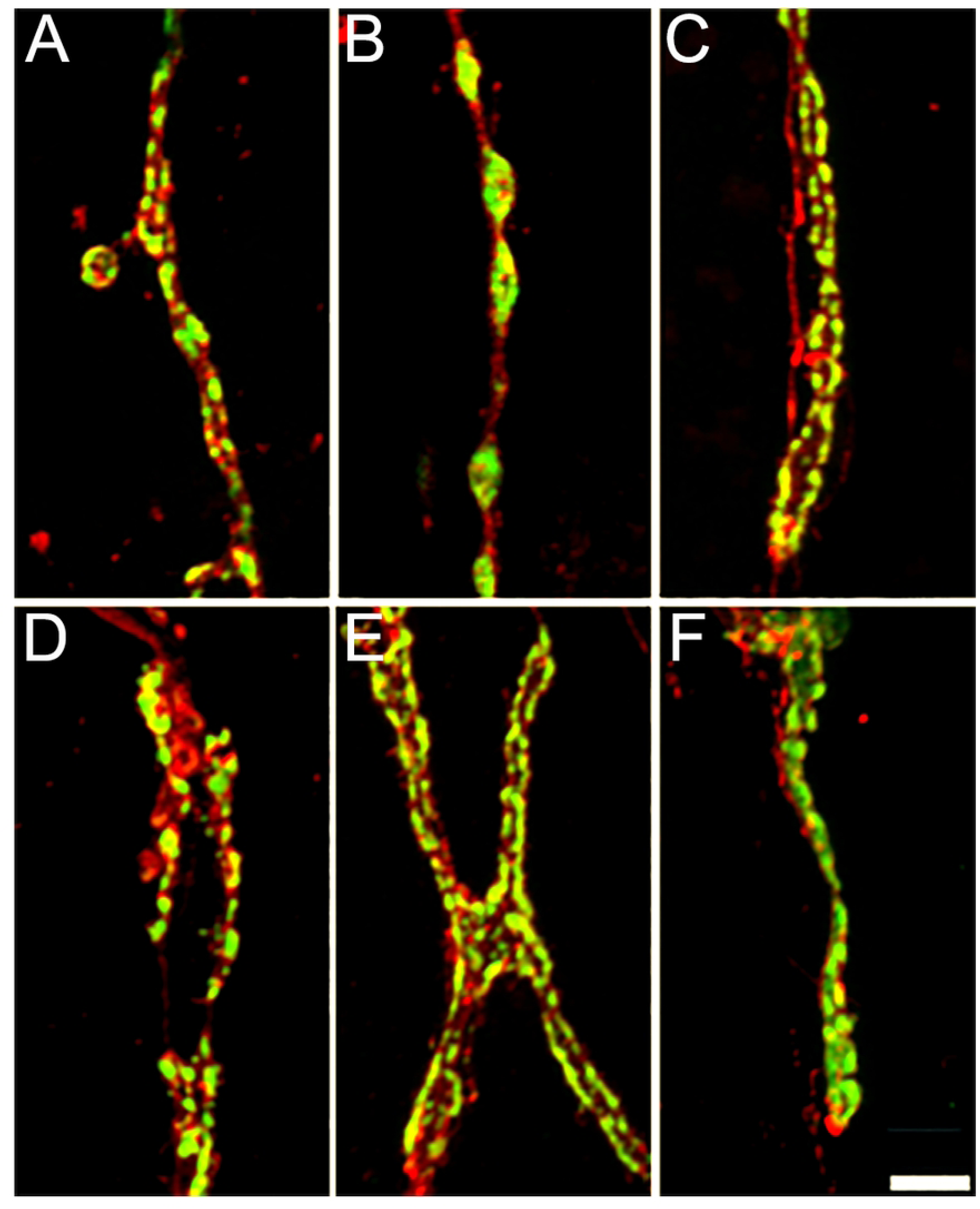
Diverse synaptic boutons structures form the NMJs of the first abdominal segment. Anti-HRP and anti-Synapsin immunoreactivity reveal different synaptic bouton morphologies at different NMJs within the first abdominal segment. For example: **(A)** large size VLI1 NMJ, **(B)** medium size VEL2 NMJ, **(C)** Small size NMJ LO1, **(D)** Very small size LT2 NMJ, **(E)** Very small size LO 4-6 NMJ, **(F)** Very small size VO1 NMJ. Scale bar: 5 µm.

NMJs are present throughout all the musculature and exhibit great variability in size and shape (Figure 5 and 6). We observed and defined four types of NMJs based on their sizes: large, medium, small, and very small (Figure 5A-D). Large size NMJs are located in muscles VIL 1 to 3 and VEL 5 and show very small varicosities that do not produce quantifiable synaptic boutons (Figure 5A and 6A). Medium size NMJs are located in muscles VEL 2 to 4 and VEL6 and show very well distributed varicosities that form quantifiable synaptic boutons (Figure 5B and 6B). Small size NMJs are located in muscles VIL 1 and LO 1 showing small varicosities that do not produce quantifiable synaptic boutons (Figure 5C and 6C). Very small synapses are located in muscles VO 1, LT 1 to 4, and LO 2 to 9 (Figure 5 and 6D-F). Synaptic terminals in LT 1 to 4 show a mix of tightly grouped and very distributed varicosities that produce quantifiable boutons of various sizes (Figure 6D). Varicosities in VO 1 and LO 2 to 9 are very condensed, producing mainly unquantifiable boutons, although quantifiable boutons can also be observed (Figure 6E-F).

### Selection and description of the model NMJ VEL 2

We selected a model NMJ based on multiple criteria: 1) the NMJ is innervating a muscle that is easy to locate, 2) is easy to differentiate from other synapses, and 3) can be easily visualized. Furthermore, we considered the fact that its size is not an obstacle for its study. Indeed, a large synapse would not be adapted to a time- and energy-efficient study, while a small synapse might not allow the future characterization of undergrown synapses. In addition, its synaptic boutons must be easy to visualize and belong to a synapse with quantifiable boutons (see previous section).

The NMJ located on muscle VEL 2 fits all our criteria. This NMJ is a medium size synaptic terminal that originates from the “ventral nerve a” tract and is located in the external ventral muscle layer of the first abdominal segment. Its typical structure has two main branches covered with synaptic boutons, but secondary branches are also commonly observed. Its synaptic boutons are rounded, well distributed and cover a great part of the muscle (Figure 7A-C). The number of synaptic boutons observed in different VEL 2 synapses is normally distributed (Shapiro-Wilk, p =0.8) and ranges between 47 to 140 (*X̅*: 86.89; σ: 24.04, n = 19; Figure 7D). The muscle innervated by this terminal ranges from 22017 to 69982 μm^2^ (*X̅*: 44627; σ: 13452, n=19; Figure 7E) and shows a normal distribution (Shapiro-Wilk, p =0.82).

**FIGURE 7:**
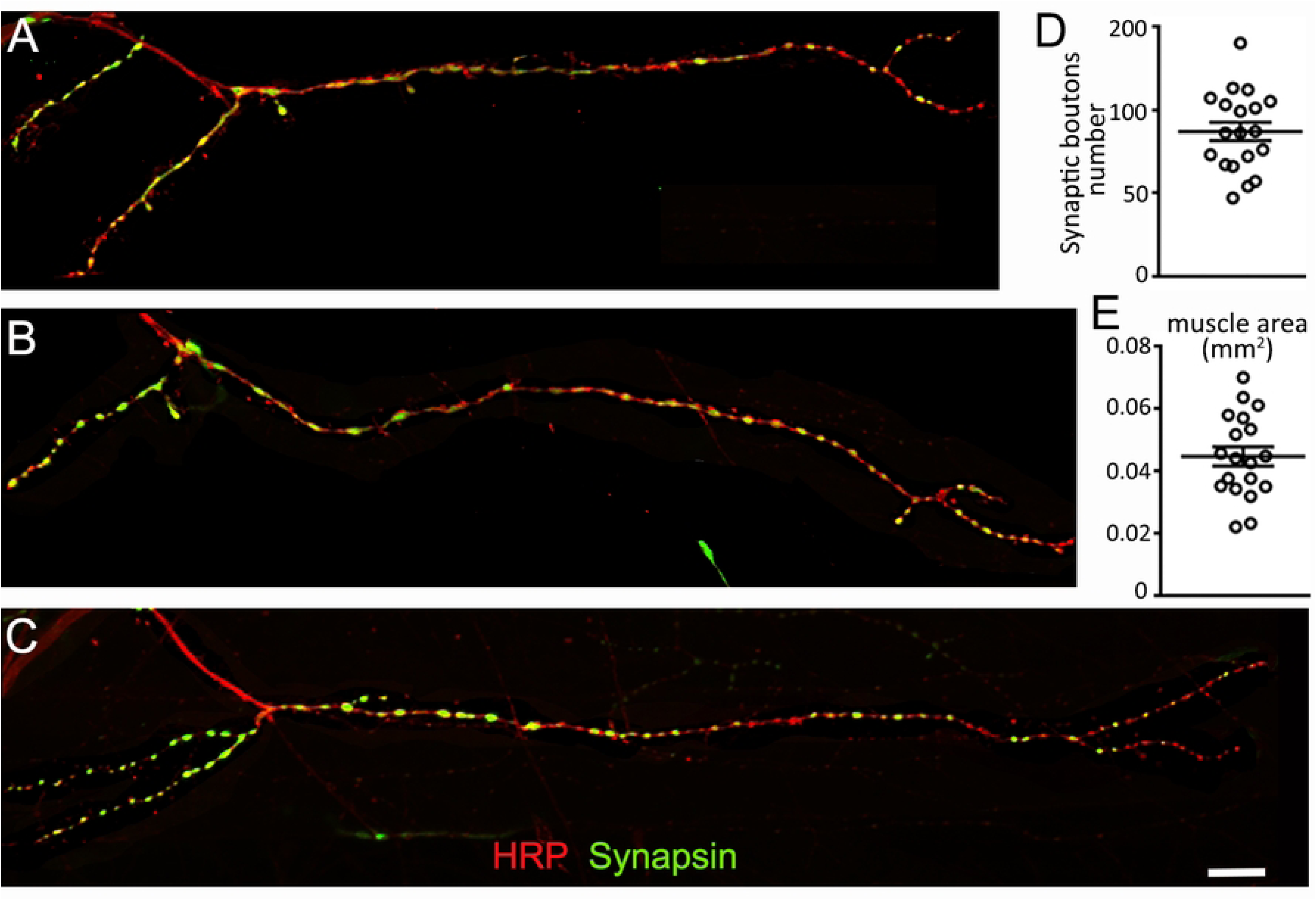
The VEL 2 model NMJ. **(A-C)** Confocal images of 3 different NMJs from muscle VEL2. Anti-HRP (red) and anti-Synapsin (green) immunofluorescence reveal the structure of the VEL2 synapse from 3 different animals. Scale is 50 µm. **(D)** Quantification of the number of synaptic boutons and **(E)** muscle size. Data is shown as a scatter plot and mean ± SEM.

In this work we have shown the basic architecture of the nervous system and muscles of *C. vitellinus*. We then focused our attention on a specific segment (Abdominal 1) and described in detail all the different muscles and their connecting nerves. Finally, we considered all the NMJs in this segment and selected a model NMJ for future study. While the inspiration for this work is the *Drosophila* NMJ, it remains to be seen whether we will be able to use the *Chironomus* NMJ to characterize changes in synaptic growth.

## DISCUSSION

The *Chironomus* larval NMJ shows great potential to be used as a cellular and tissue marker for future investigative work assessing rivers contamination. Indeed, *Chironomus* larvae inhabit river sediments (site of contamination) and work with *Drosophila* has shown that the NMJ responds to external/environmental factors (e.g., Sigrist et al., 2003; Zhong and Chun-Fang Wu, 2004; and Hirsch et al., 2012). A description of the *C. vitellinus* neuromuscular system was therefore a necessary preliminary work.

Although partially known, the *Chironomus* neuromuscular anatomy has always been incomplete. Miall and Hammond’s work on *Chironomus dorsalis* presented a general view of the system, only describing the first layers of muscles without naming them and relating them to the nervous system [15]. Nerve tracts and the NMJs were not described by these authors nor by recent studies describing the nervous system in other *Chironomus* species [6,8,18]. The description presented in this work attempts to remediate these issues and presents a detailed anatomical description that could be used for toxicological studies.

The *Chironomus* larval abdominal muscular anatomy presented in this work is the most detailed available. It is unique when compared with the anatomy of other abdominal muscle arrangements described in other insect species such as *D. melanogaster* [29], *Periplaneta americana* [45], and *Galleria mellonella* [46]. Comparing the muscular anatomy of *Chironomus* with other insect anatomies is challenging since current insect descriptions use different body sections to characterize the muscle distribution. In this study, we focused on describing the first abdominal segment since the abdominal region of the larval stage is the most used for movement. Similar abdominal descriptions are common for the crawling larvae of insects (e.g., *D. melanogaster* [30]). Nevertheless, studies describing the anatomy of adult winged insects typically examine the thoracic section of the body given its relation to movement (e.g., *Hemianax papuensis* [47]). It is also true for the musculature descriptions of walking insects with well-developed appendages; in this case, the descriptions of the thoracic and appendage musculature are preferred (e.g., *Locusta migratoria*[48]).

When comparing the *Chironomus* anatomy to other crawling larvae such as *D. melanogaster,* we can notice that some muscle arrangements in the first abdominal segment are preserved while others are completely missing. This is the case for the *D. melanogaster* ventral acute and ventral transverse muscles, which are completely missing in *Chironomus*. In contrast, the lateral transverse and segment border muscles are preserved in both species. Another singular feature of the *Chironomus* musculature is that there are multiple lateral oblique (LO) muscles compared to the single LO muscle structure observed in *D. melanogaster*. The same is observed with the ventral oblique (VO) muscles, but this time the *Chironomus* anatomy only shows a single VO muscle structure while *D. melanogaster* shows multiple. The number of muscles per hemisegment in the first abdominal segment of the studied *Chironomus* species is similar to the number of muscles in *D. melanogaster* (31/*Chironomus* hemisegment vs 30/ *D. melanogaster* hemisegment [49] however, their distributions through the ventral, lateral and dorsal regions differ between species.

The basic structure of the larval central nervous system was known in other *Chironomus* species [6,8,15] and *C. vitellinus* shows similar structure. We did, however, characterize a number of new features such as the different nerve tracts originating from the main nerve and their multiple NMJs. Nerve tracts trajectories in *Chironomus* bear similarities to the trajectories of the nerve tracts observed in *D. melanogaster*. The *Chironomus* dorsal nerve trajectory appears to be homologous with the *Drosophila* intersegmental nerve (ISN) trajectory, the *Chironomus* ventral a-c nerve trajectory to the *Drosophila* segmental nerves b-d (SNb-d) and the *Chironomus* lateral nerve trajectory to the *Drosophila* segmental nerve a (SNa). Nevertheless, tracing techniques such as retrograde labeling with different dyes (Inal et al., 2020) will need to be performed to corroborate the exact nerve trajectories in the abdominal segment of the *Chironomus* larvae. Pursuing this method should also allow the identification of the motor neuron cell bodies innervating the muscles.

Synaptic terminals in the *Chironomus* tissue show a large variability of sizes paralleling the high variability in muscle size observed through the larval tissue. The great variability of synaptic varicosities made it difficult to identify suitable NMJs with quantifiable synaptic boutons. Similar variations in varicosities were previously observed in *Chironomus tentans* when trying to detect Bombesin immunoreactivity in nerve fibers of the larval alimentary canal [17]. The quantifiable NMJs in the ventral and lateral area can only be identified in the ventral external longitudinal (VEL) muscles. These muscles present an assortment of rounded and well-formed synaptic boutons making their quantification easy under the microscope. NMJs on lateral transverse (LT) muscles also present quantifiable synaptic boutons, but their quantification is difficult due to the presence of large amounts of fat tissue obstructing these terminals.

The model NMJ (VEL2) was selected due to its visible location in the tissue and due to its quantifiable boutons. It is characterized by its large size, separated bouton arrangement and simple structure with elongated branches. The number of synaptic boutons of the *Chironomus* model NMJ show similarities with the number of synaptic boutons observed in *Drosophila* at the model 6/7 NMJ. Authors have reported values averaging 75 to 100 boutons per NMJ in wild type *Drosophila* larvae [50–52]. For example, in our laboratory, a recent analysis of the Drosophila 6/7 NMJ at segment A3 shows a mean of 79 boutons when we examined 19 synapses (data not shown). Our data ranged from synapses containing 48 boutons to synapses containing 116 boutons and the sample shows a standard deviation of 16. The *Chironomus* model NMJ shows a mean average of 87 synaptic boutons for synapses ranging between 47 and 140 (standard deviation 24; n = 19; Figure 7D). The variability observed in *Drosophila* studies is lower than the ones observed in *Chironomus.* This is easily explained by the fact that the *Drosophila* strains are isogenic while the studied *Chironomus* animals are the progeny of genetically varied animals collected in the field. This observed variability in the number of boutons could also be due to factors such as larval size, sex, and age. Some of these factors are known to affect the structure and function of synaptic terminals in *Drosophila* [53,54] thus similar effects could be occurring in the *Chironomus* NMJ. Future experiments will have to be undertaken to assess whether these factors play a role. If they are, it should be possible to reduce the *Chironomus* synaptic size variability by only considering a well-defined subset of animals.

The present study of the *Chironomus* neuromuscular system provides the foundation upon which further research could develop. It identifies basic NMJ features that could be used as markers for assessing toxicity. In addition to quantifying the number of boutons and the muscle area, other features of the synapse (span, arborization, active zone numbers, postsynaptic differentiation) could be analyzed and quantified as new immunohistochemical tools become available. Given the outstanding neuromuscular anatomy of *Chironomus*, procedures such as electrophysiology could also be used to determine disruptions on neuronal signaling and synaptic transmission caused by toxic chemicals. Due to the versatility of this system, we are hopeful that it could be used to evaluate the responsiveness of the NMJ to biological, environmental, and toxicological variables.

## ACKNOWLEDGEMENTS

This work was supported by the Puerto Rico Center for Environmental Neuroscience (PRCEN) under NSF grant-HRD 1736019 and by the NIH-NIGMS grant GM103642 to Bruno Marie. Infrastructure support and service were provided, in part, by the grant U54MD007600 from the National Institute on Minority Health and Health disparities (NIMHD). Monoclonal antibodies were obtained from the Developmental Studies Hybridoma Bank, created by the NICHD of the NIH and maintained at The University of Iowa, 45 Department of Biology, Iowa City, IA 52242. We would like to acknowledge Dr. Budnik and Dr. Di Antonio for the gift of antibodies (rabbit anti-Dlg and rabbit anti-GluRIII). We thank Dr. Blagburn for his valuable comments on previous versions of this manuscript.

